# Semi-automated image analysis of root architecture and early root development in faba bean and white clover and genomic estimation of breeding values and correlations

**DOI:** 10.64898/2025.12.01.691692

**Authors:** Istvan Nagy, Peter Skov Kristensen, Elesandro Bornhofen, Marta Malinowska, Linda Kærgaard Nielsen, Andrea Schiemann, Niels Rolund, Stig Uggerhøj Andersen, Torben Asp

## Abstract

Protein-rich leguminous plants, such as faba bean and white clover are prospectively interesting crops in the North-European countries for reducing dependence on soybean import. Significant expansion of the production area of leguminous crops is challenged by the sub-optimal climatic conditions in this region, especially by the increasing probability of year-to-year fluctuation of extreme weather conditions due to global climate change. To overcome these challenges, development of new climate-resilient varieties suitable for growing under Northern-European conditions are needed. Root architecture and early root development, as well as the availability of efficient root phenotyping technologies are crucial factors of advancing in breeding of adequate varieties. We report a study of a simple and affordable screening technology of early root development using rhizoboxes in connection with semi-automated image analysis and provide a conceptual pipeline for estimation of Genomic Estimated Breeding Values (GEBVs) and correlating greenhouse and field phenotype data. Based on bivariate models, high genetic correlation (r=0.83) could be detected between total root length values recorded in greenhouse rhizobox experiments and field grain yield in faba bean. In white clover, moderately positive genetic correlation (r=0.17) between estimated breeding values of rhizobox-detected total root length and field yield could be identified. Our results suggest that phenotyping and selection of early root development components could potentially be useful in breeding programs to increase the genetic gain for field yield.

## Introduction

Agriculture and food industry of Northern European countries, including Denmark are greatly depending on import of crop products providing protein resources - mainly soybean and processed soybean products. Denmark imports about 1.7 million tonnes of soybean meal annually (approximately 0.7% of global soy production), primarily for livestock feed.^1^ To reduce dependence on soybean import, there is an increasing interest in the Nordic countries in expanding local production of protein-rich leguminous crops, like faba bean (*Vicia faba* L.) and white clover (*Trifolium repens* L.).^2^

Expansion of the area of leguminous crops is beneficial for environmental-friendly agriculture, as legumes can fix atmospheric nitrogen, reducing the input of chemical fertilizers.^3^ Notably, white clover can fix up 270 kg nitrogen per hectare through association with *Rhizobium leguminosarum* bv. *trifolii*.^4^ and the nitrogen fixing capacity of faba bean is estimated up to 160 kg nitrogen per hectare, of which about 50% will remain in the soil after harvesting. ^5^

Legumes fit well into the crop rotation, alternating with cereals, tubers or industrial crops, like rapeseed or sugarbeet, because the spacious legume root system remediates soil structure by creating stable biopores in the subsoil, improving water infiltration, and root penetration for subsequent crops.^6^ In addition, the deep root system of legumes contribute to long-term soil carbon storage by stabilizing organic carbon below the frequently disturbed topsoil. ^7^

The most limiting factors of extending the leguminous protein crop production area in the Nordic region are the sub-optimal climatic conditions, particularly the increasing probability of year-to-year fluctuation of extreme weather conditions driven by global climate change. Projections and models show that in Northern Europe the trend of increased winter precipitation and decreased summer precipitation and the intensifying of summer heatwaves will continue.^8^ Spring drought episodes can critically restrict water availability during reproductive stages, while water abundance in cold winters may lead to temporary soil saturation and subsoil hypoxia, affecting root development and resource uptake.^9,10^ These challenges underline the need of developing new climate-resilient varieties suitable for growing under Northern-European conditions.^11^ Root architecture and early root development (i.e., root development dynamics during the first few weeks after germination), together with the availability of efficient root phenotyping technologies are therefore important factors for advancing breeding of adequate varieties.

Root System Architecture (RSA) increasingly attracts the interest of scientist in the era of climate change. RSA comprises the shape and spatial arrangement of the root system within the soil ^12^ and is determined by genetic-as well as by environmental factors, like soil type, soil temperature and supply of water and nutrients, while roots exhibit a high level of phenotypic plasticity in response of environmental conditions.^13^ Evidently, root architecture is a key factor of efficient nutrient and water uptake, and even a ‘Second Green Revolution’ has been envisioned that deploys crops with improved below ground traits.^14,15^ The development of new plant varieties with improved root phenotypes requires advances in the characterization of root development and morphology, as well as in the relationships between genetic factors and RSA. As being a complex, polygenic trait, studying of RSA should be conducted by parameterization of its elements, such as root depth, total root length, root angle, root thickness, root density and root surface area.^16,17^

A wide range of root phenotyping technologies have been developed and applied during the last decades, ranging from excavating root crowns on the fields (“shovelomics”^18^) to semi-field based micro-rhizothron facilities with automated image acquisition. ^19^ Non-destructive 3D root phenotyping using tomographic techniques, like X-ray computed tomography, magnetic resonance imaging (MRI) and positron emission tomography (PET) also become achievable during the last decades (refer^15^ for a review).

Soil-filled slab chambers with transparent lids (rhizoboxes) offer a reasonable and space-efficient approach for studying early root development under controlled conditions. Rhizoboxes allow the visualization and non-invasive quantification of early root development *in situ*, as they does not disturb the spatial disposition of roots. Although they diminish the root system’s growing space in an artificial substrate, thereby quantitative data obtained from rhizobox experiments not directly refer to root development under field conditions, but robust rhizobox results obtained under controlled conditions can nevertheless be extrapolated to form a better understanding of expected root development in the field, or to identify particular traits of interest in breeding. ^20^

Investigations on early root development are of fundamental importance, as a vigorous early root system is essential for healthy plant growth and establishment, especially under sub-optimal conditions,^21^ and recent quantitative genetics studies in perennial ryegrass demonstrated that early root development traits can act as proxies for field yield across multiple years and locations. ^22^ Earlier studies identified correlations between early root traits and crop productivity and/or nutrient- and water use efficiency in other species, like maize^23,24^ and rice^25^ as well. Such relations have been confirmed by the co-occurrence of quantitative trait loci (QTL) between early root traits and yield components in maize- and cereal field experiments.^24,26,27^ On the other hand, to date limited information is available concerning correlations of early root development and field yield components in legumes.

The decreasing genotyping costs and the availability of high-quality genomic references provide opportunities for breeders to obtain abundant genotyping information at affordable costs and to utilize large-scale genotype data for genomic predictions along with phenotpype data collected by multiple technologies across multiple years and locations. However, for the successful implementation of such efforts, the development of appropriate statistical models and methodologies is necessary. For polygenic quantitative traits, like RSA components, Genomic Selection (GS) offers cost effective opportunities for breeders to increase the rate of genetic gain, by reducing the time needed to complete a breeding cycle, increasing the selection intensity and by utilizing within-family variation that can be captured using molecular markers.^28^ GS is a form of Marker Assisted Selection that simultaneously estimates all locus-, haplotype- or marker effects across the entire genome to calculate Genomic Estimated Breeding Values (GEBVs^29^). Multivariate models are valuable for integrating multiple correlated traits into the breeding process, and estimation of genetic correlations between traits and can potentially improve the accuracy of GEBVs.^30–32^

Yield is often an important breeding goal, but it is relatively expensive and time-consuming to get phenotypic observations for yield, since varieties ideally should be tested in replicated field trials in several locations and several years due to potential spatial variation within fields, and due to Genotype by Environment (GxE) interactions across different environments, which complicates the selection of genotypes consistently outperforming under variable environmental conditions. Yield cannot be quantified directly and accurately in early phases of breeding programs, when only one or few plants are available per variety. However, if other traits can be identified that are genetically correlated to yield and that can be phenotyped more easily and/or in shorter time in controlled conditions, using a few seeds or plants per variety, then such traits would be useful to implement in early selection stages of breeding programs in order to increase the genetic gain of yield.

To facilitate quantitative genetics studies for correlating early root development traits and yield components, we conducted greenhouse rhizobox experiments using faba bean and white clover genotypes, including standard varieties and breeding lines. The specific objectives of the study were to quantify early RSA traits using image-based phenotyping, assess genetic variation and heritabilities using genomic data, and evaluate genetic correlations between early RSA traits and agronomic performance to determine the potential of early root phenotyping under semi-controlled conditions for indirect selection in breeding programs.

## Materials and methods

### Plant material

For faba bean, 180 spring lines from the breeding program of the Danish breeding companies Nordic Seeds (75 lines) and Sejet (75 lines), as well as 10 standards varieties and 20 core lines belonging to the IMFABA Consortium (https://projects.au.dk/fabagenome/genomics-data) were used in the study (Table S2). For white clover, a panel consisting of 175 lines selected from 20 commercial varieties with diverse agronomic characters was used (Table S3).

### Rhizobox setup and growing conditions

Plants were grown in custom-made plastic rhizoboxes (internal dimensions: 36×18×2.5cm). Boxes were filled with 1.8L substrate, containing 85% (v/v) turf-based substrate (Hawita Uni20, Hawita GmbH, Germany), 10% (v/v) local topsoil and 5% (v/v) sand (washed and fractionated, 1.4 to 2.5mm), and supplemented with 300ml water. In case of faba bean, two seeds were sown in each box. In order to minimize within-line genetic diversity, initial white clover plant material was maintained in the greenhouse as clonally propagated stocks in 17 cm pots. For white clover rhizobox experiments, 8 to 12 cm long cuttings of young lateral shoots were collected from the middle part of the pots. It was attempted that cuttings have a nodus above the cutting site and fully developed leaflets below the shoot tip. During processing, cuttings were kept between wet filter paper sheets. Boxes were placed in the greenhouse on racks that ensured to incline them at 60° with backward-leaning transparent side, to facilitate root growth along the visible side. During growing, the transparent plexiglass front side of the boxes was covered with a black foil. For both plant species, experiments were conducted in three subsequent batches in randomized blocks throughout a 12 month period in the same greenhouse compartment equipped with an automatic shadowing system during summer and additional lighting and heating (16/8h day/night at 21/18 °C) during winter and without an active cooling system in summer. In each replicate, plants were grown for 25 to 28 days, by applying manual watering with equal amounts (100 to 200 ml, depending on humidity and outer temperature) of tap water on every second day, applied over the full soil surface of rhizoboxes.

### Phenotype data collection

#### Root imaging

On the final day of growing period, in each batch experiment root images were taken at 600dpi resolution by a photo quality scanner (Epson Perfection V700). The built-in (removable) lid of the scanner was replaced by a custom-made framed glass plate, covered by a non-transparent black fabric, allowing to scan rhizoboxes in horizontal position. Root imaging was conducted by the RootPainter software. ^33^ The analysis procedure involved training with an initial manual annotation and iterative corrective annotations in order to get a refined segmentation model, which was used to automatically segment the entire root dataset. Finally, RootPainter was used to extract Total Root Length (TRL) values for each segmented image. In addition, segmented root images were exported and further processed by the RhizoVision software^34^ to quantify and record the following root phenotype components: Average root diameter, Number of Branching points and Root surface area. (Find details of root imaging in the Supporting information.)

#### White clover leaf phenotyping

For leaf phenotyping, in each genotype, three fully developed trifoliate leaves were collected from the source plants maintained in the greenhouse. Separated leaflets (9 single leaflets of three trifoliate leaves for each genotype) were fixed on white paper sheets by slight glueing their back side and scanned in full-color mode at the highest possible resolution. The scanned images were used for digital image analysis to determine leaf size, leaf shape and leaf color. Leaf size (average surface area of leaflets) and leaf shape parameters (circularity and solidity) were determined by ImageJ (v1.54g).^35^ For leaf color analysis, 20×20 pixel squares were cut out from the middle part of the leaflet images and saved to separate files using the image editing software Inkscape.^36^ Average pixel RGB values were calculated from the 400 px image files by the *convert* tool of the ImageMagick software^37^ and for each line, RGB values were converted to perceived brightness values (Formulae S1, S2 and S3).

#### Faba bean field phenotype data

Faba bean breeding lines and variety standards were tested in replicated multi-year field trials in three years (2022, 2023, 2024) at three locations (Dyngby, Svinø and Gamborg, Denmark) in alpha-lattice design with 10 m^2^ plots in three replicates. Plant height, grain yield and Thousand Kernel Weight (TKW) were recorded. Moisture and protein content was determined from harvested seeds using two rapid, non-destructive optical technologies, Near-Infrared Transmission (NIT) ^38^ and Near-Infrared Reflectance (NIR),^39^ resulting in ca. 4000 observations in total for each trait.

#### White clover field phenotype data

A two year randomized field plot experiment was conducted in 2021 to 2022 in Haldrup, Denmark, with 12.5 m^2^ plots in two replicates. Fresh and dry biomass yield was recorded after four cuts in the first year and five cuts in the second year.

#### White clover greenhouse biomass data

Image-based biomass measurements were carried out on a panel of white clover lines overlapping with the panels of rhizobox- and field experiments. For each line, 10 individual plants were monitored using stationary cameras to estimate cumulated above-ground biomass after 10 days of growth.^40^

### Genetic variant data

#### Faba bean variant data

Genetic variant data based on pseudo-chromosome references of the latest version of the *Vicia faba* genome sequences (Hedin/2 v.2), ^41^ containing 122,291 bi-allelic Single Nucletide Polymorhisms (SNPs) was provided by the IMFABA consortium (https://projects.au.dk/fabagenome/genomics-data). Of the 180 lines used for the rhizobox experiments, 157 were represented in the VCF file. Based on the variant data stored in the VCF file, three-dimensional Prinicipal Component Analysis (PCA) plots were generated using VCF2PCACluster. ^42^

#### White clover variant data

Genetic variant data based on Genotyping by Sequencing (GBS) short-read sequences mapped onto the *Trifolium repens* v.4.0 genome references. ^43^ Of the 195 genotypes represented in the original variant discovery project, 167 were common with the line set used in the root phenotyping experiments, therefore variant data for the not matching lines were removed from the VCF file and genomic analysis studies was further on restricted to these 167 lines. The VCF file was further on filtered for Minor allele frequency >3%, Number of missing data <30% across lines and a minimum Read Depth of 5, keeping bi-allelic variants on 280,265 loci.

### Genomic analysis conception

#### Calculating variance components and heritabilities

Genomic Best Linear Unbiased Prediction (GBLUP) models were used in order to partition environmental, spatial, and genetic effects from each other for the different traits. Here, genomic relationships between lines based on SNP markers were included in order to estimate breeding values (additive genetic effects) for each trait, heritabilities (i.e. the ratio between genetic variance and total phenotypic variance), and to estimate the genetic correlations between traits. ^44^

Variance components of all models were estimated by restricted maximum likelihood using the software package DMU for analyzing Multivariate Mixed Models (provided by the Center of Quantitative Genetis and Genomics, Aarhus University ^45^).

Heritabilities at field plot or rhizobox level were calculated as narrow sense using the following formula:

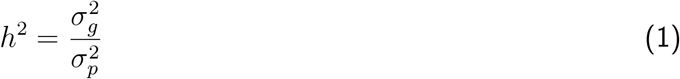

and as broad sense using the following formula:

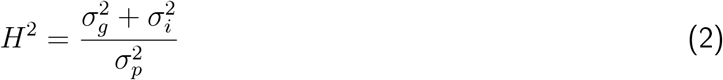

where 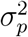 is the sum of all estimated variance components within each of the models, 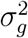 and 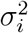 are variances of additive genetics and of line effects, respectively.

Correlations between traits were calculated as Pearson correlations between estimated additive genomic breeding values for the respective traits.

#### Correlated response to selection

The genetic response to selection per generation of a trait were calculated using the breeder’s equation:

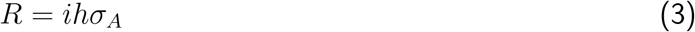

where *i* is selection intensity, *h* is the square root of the narrow-sense heritability, *σ*_*A*_ is the genetic standard deviation of the trait (i.e. the square root of the genetic variance). The genetic gain of a trait can be improved by incorporating information from another genetically correlated trait. To determine how much genetic gain can be achieved for a primary trait *(Y)*, when selection is made based on a secondary trait *(X)*, the correlated response to selection can be calculated as:

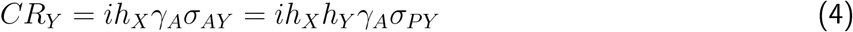

where *h*_*X*_*h*_*Y*_ *γ*_*A*_ is the co-heritability of the two traits *X* and *Y, γ*_*A*_ is the genetic correlation between traits *X* and *Y*, the genetic and phenotypic standard deviations of trait *Y* are *σ*_*AY*_,*σ*_*AP*_ and *σ*_*PY*_ respectively, according to Falconer and Mackay. ^44^

Thus, the genetic gain of trait *Y*, when selecting indirectly based on trait *X*, will better in cases where the heritability of trait *X* is high, where the genetic correlation between the traits is high, or where the selection intensity can be increased.

Dominance and epistatic genomic relationships were not explicitly included in the models, since large data sets are required to estimate such non-additive genetic effects. Instead, the line variance was included in order to capture non-additive genetic variance as well as any additive genetic variance that was not captured by the SNP markers used in the genomic relationship matrix. The line variance can only be estimated for the traits, where phenotypic observations are available for more than one replicate, since the line variance will be confounded with the residual variance if only one observation is available per line.

#### Verifying genetic correlations

For entry means of two correlated traits (for example total root length *vs*. grain yield), a simple bivariate Bayesian model was fitted using the R package *brms*.^46^ Each response included a random intercept for entry, with residual correlations fixed to zero. Models were run for 10,000 iterations (1,000 warm-up; thinning = 5; one chain)

## Models

### Faba bean models

**Yield data from field trials was modelled by:**

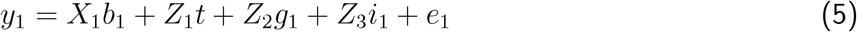

where *y*_1_ is a vector of phenotypic observations, *X*_1_ is a design matrix for fixed effects, *b*_1_ is a vector of effects of year-location-replicate, *Z*_1_, *Z*_2_ and *Z*_3_ are design matrices for random effects, *t* is a vector of effects of blocks within year-location-replicate with 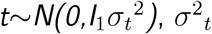 is the variance of block effects, *g*_1_ is a vector of additive genomic breeding values with 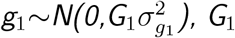 is a genomic relationship matrix calculated using method one proposed by VanRaden, ^47^ 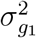 is the additive genetic variance, *i*_1_ is a vector of line effects with 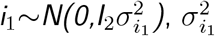 is the variance of line effects, and *e*_1_ is a vector of residual effects with 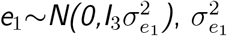 is the variance of residual effects, and *I*_*n*_ are identity matrices.

**Root length data from rhizoboxes was modelled by:**

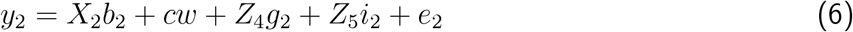

where *y*_2_ is a vector of phenotypic observations, *X*_2_ is a design matrix for fixed effects, *b*_2_ is a vector of effects of experiment, *c* is a fixed regression coefficient for observed total root lengths regressed on thousand seed weights, and *w* is a vector of Thousand Kernel Weights (TKW) of seeds sown in rhizoboxes, *Z*_4_ and *Z*_5_ are design matrices for random effects, *g*_2_ is a vector of additive genomic breeding values with 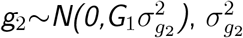 is the additive genetic variance, *i*_2_ is a vector of line effects with 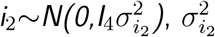 is the variance of line effects, and *e*_2_ is a vector of residual effects with 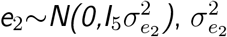 is the variance of residual effects, and *I*_*n*_ are identity matrices.

Combining the two models above, a bivariate model was used for estimating genetic covariances between field yield and rhizobox root length traits:

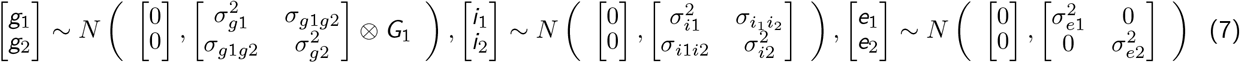

The remaining random effects were assumed normally and independently distributed with zero means and variances as described above.

The genetic correlation between traits, 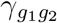, can be calculated from the estimated genetic variances and co-variances:

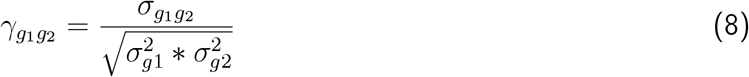

### Clover models

**Yield data from field trials was modelled by:**

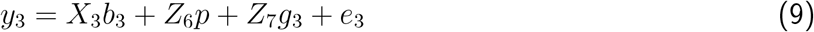

where *y*_3_ is a vector of phenotypic observations, *X*_3_ is a design matrix for fixed effects, *b*_3_ is a vector of effects of cut number, *Z*_6_ and *Z*_7_ are design matrices for random effects, *p* is a vector of effects of plot with 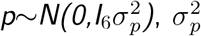 is the variance of plot effects, *g*_1_ is a vector of additive genomic breeding values with 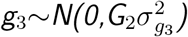, *G*_2_ is a genomic relationship matrix, 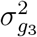 is the additive genetic variance, and *e*_3_ is a vector of residual effects with 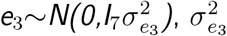 is the variance of residual effects, and *I*_*n*_ are identity matrices.

**Data from greenhouse was modelled by:**

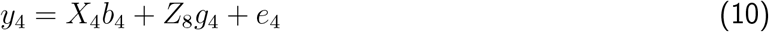

where *y*_4_ is a vector of phenotypic observations, *X*_4_ is a design matrix for fixed effects, *b*_4_ is a vector of effects of experiment, *Z*_8_ is a design matrix for random effects, *g*_4_ is a vector of additive genomic breeding values with 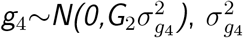 is the additive genetic variance, and *e*_4_ is a vector of residual effects with is the variance of residual effects with 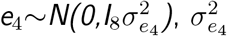 is the variance of residual effects, and *I*_*n*_ identity matrices.

## Results and discussion

### Rhizobox experiments

Rhizobox setups differed between the two species: Faba bean seeds were sown directly, while white clover plants were raised from shoot cuttings. The latter approach proved more challenging, as the greenhouse stock plants showed some inherent variability, which affected the uniformity of starting material. Due to the use of this type as starting material for white clover, the rhizobox experiments in this species targeted the re-rooting ability of explants, in contrast to the seedling root development in faba bean.This approach reflected white clover root development under natural conditions, as in this species the initial central taproot dies as the plant matures and is replaced by a secondary root system made of adventitious roots forming at each node.^48^ As the three experimental batches were conducted sequentially over a full year in the same greenhouse compartment, seasonal differences and variable greenhouse conditions (Figure S1) certainly influenced the consistency of data obtained from different experiment batches. Roots were scanned when they reached about two-thirds of the rhizobox hight: after 12 to 15 days in case of faba bean, and 15 to 20 days in case of white clover (Figure 1). RootPainter provided a reliable root image segmentation across images. Image-based quantification resulted in approximately normal data distributions for most parameters in case of faba bean (Figure 2), while in white clover the distributions were less explicit (Figure S2). Among the parameters extracted, total root length (TRL) was consistently the most robust trait, with lower sensitivity to background noise or overlapping roots. In contrast, vertical measurements, such as root diameter or perimeter were more prone to errors, as overlapping side roots frequently interfered with accurate detection. In agreement with earlier findings, ^22^ these issues underline that TRL is an appropriate RSA descriptor for downstream analyses, while caution is needed when interpreting vertical or diameter-based traits.

**Figure 1:**
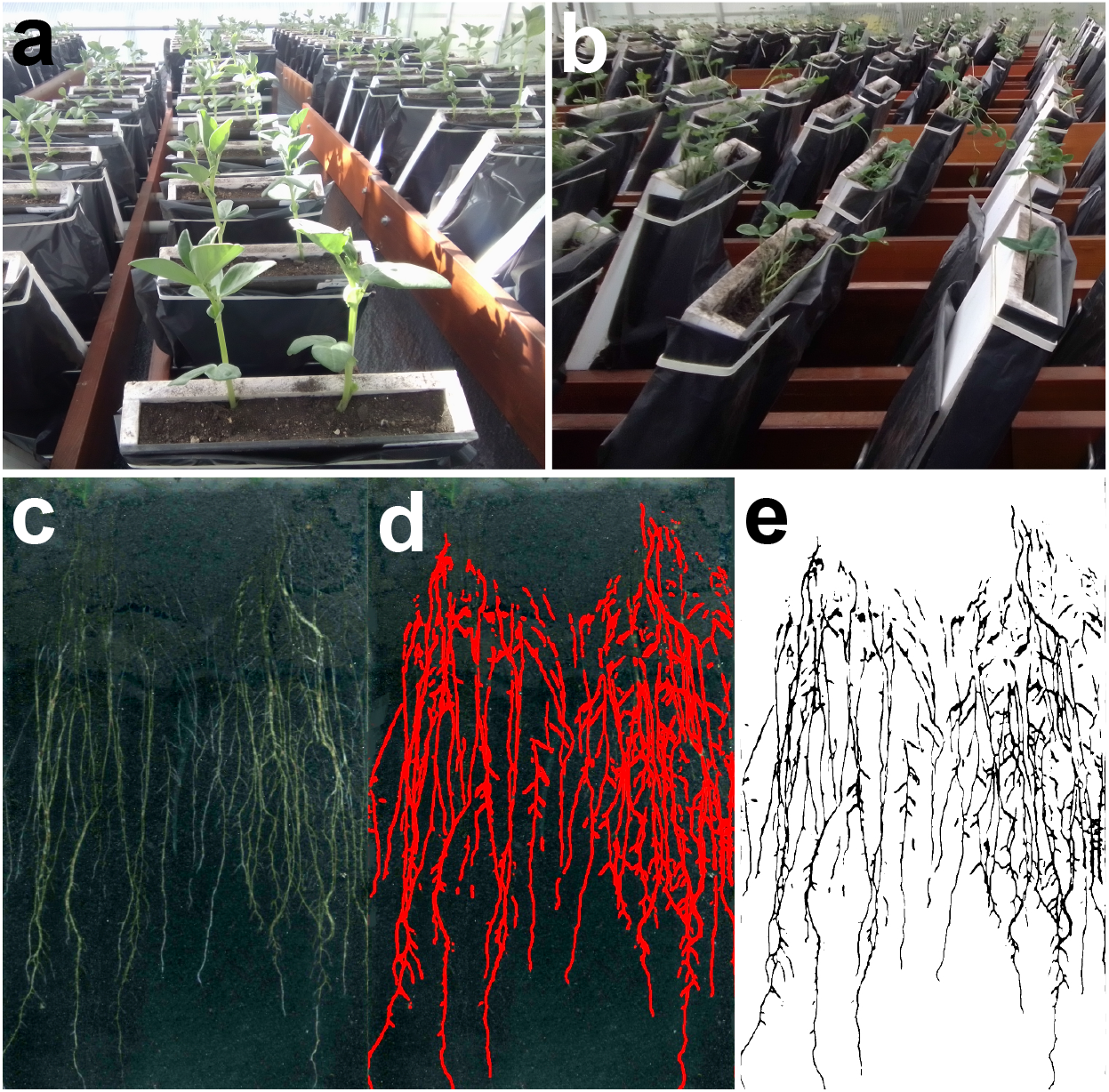
Greenhouse rhizobox experiments and root image processing. **(a)** Faba bean seedlings in rhizoboxes **(b)** White clover explants in rhizoboxes **(c)** Scanned root image of a faba bean sample **(d)** Segmentation image made by RootPainter **(e)** Segmentation image exported to analysis by RhizoVision

**Figure 2:**
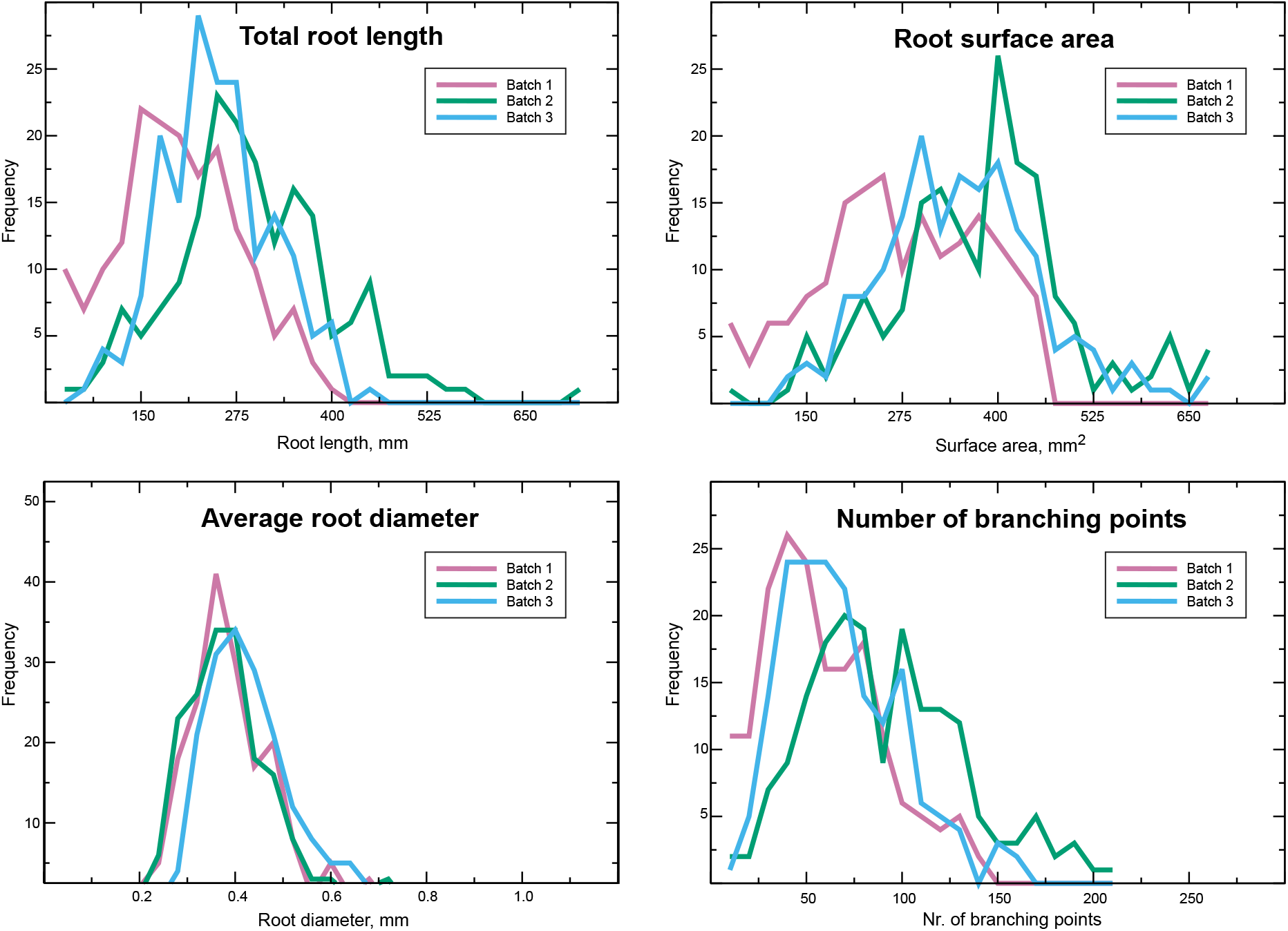
Distribution of quantitative root parameter values in three faba bean rhizobox batch experiments

### Genetic variant data, diversity and population structure

In case of both species, high-quality reference genomes and large sets of genetic variant data were available for the lines included in the study (bi-allelic SNPs on 122,291 loci for faba bean and on 280,265 loci for white clover). In faba bean, the core plant material included 75 - 75 breeding lines from two Danish breeding companies. According to the PCA plot based on genetic variant data is showing that the majority of these faba breeding lines are separating to distinct clusters according to company origin, suggesting that the breeding programs of the companies is predominantly based in different germplasm resources (Figure 3). For white clover, a panel consisting of 174 lines selected from 20 commercial varieties (4 to 10 lines for each variety) was used. In both species, the available genetic variant data enabled to build robust and reliable Genomic Relationship Matrices for genomic estimation of breeding values and correlations (Figure 4).

**Figure 3:**
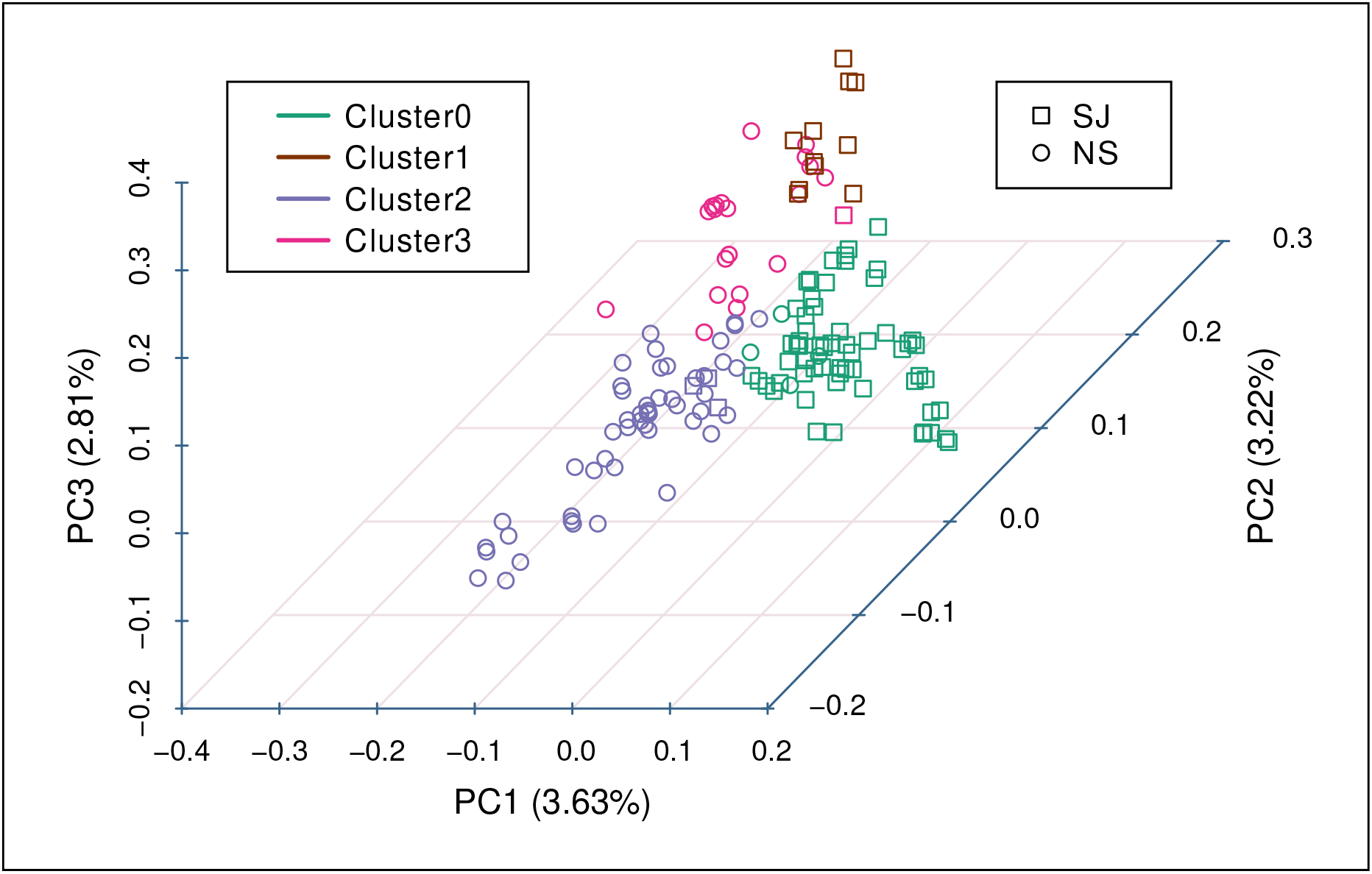
3D clustered PCA results of Faba bean breeding lines based on bi-allelic SNP markers. **SJ** and **NS** are for the Danish breeding companies Sejet and Nordic Seed

**Figure 4:**
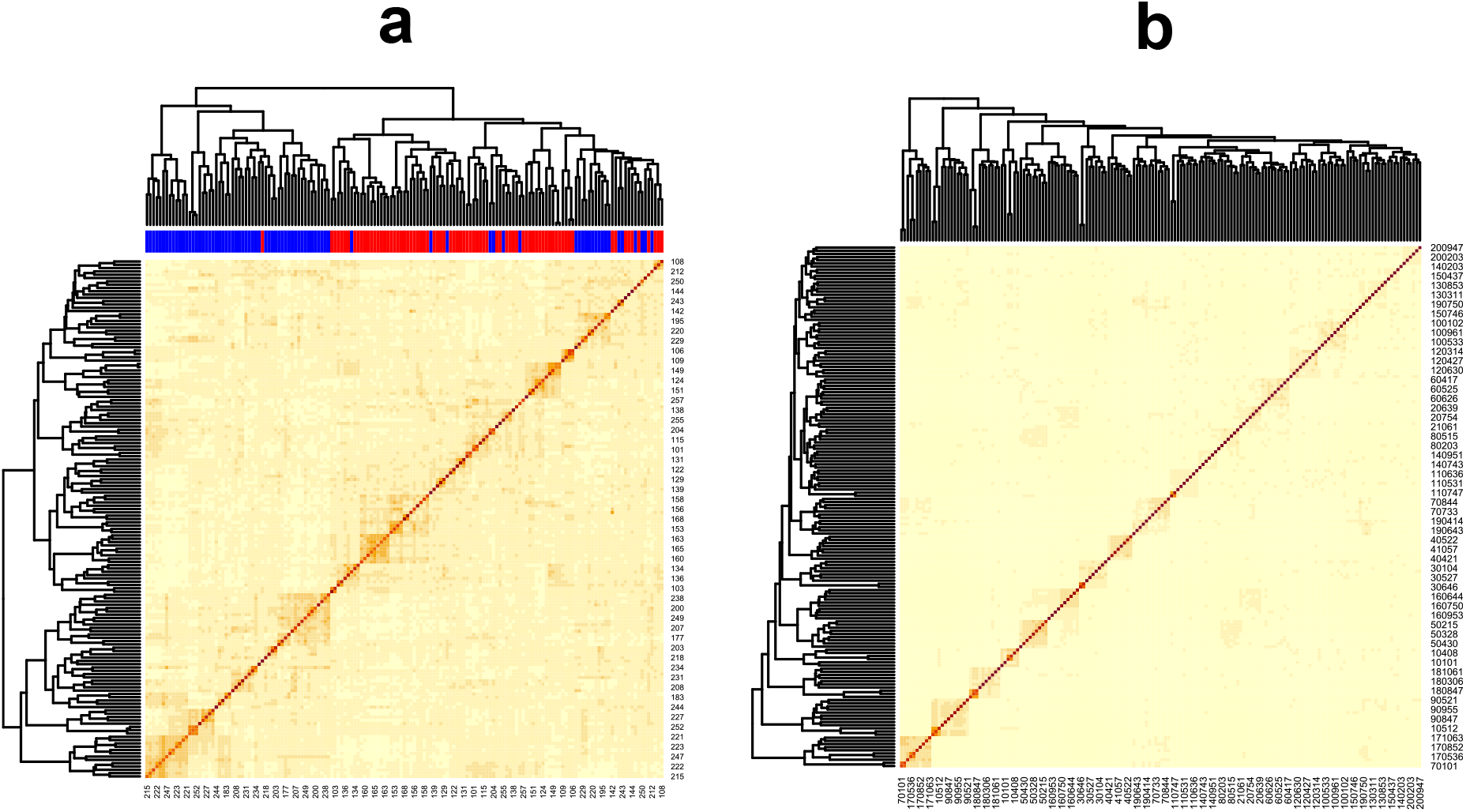
Genomic realationship matrices based on bi-allalic SNP markers. **(a)** Faba bean lines, Blue bars: **Nordic Seed** lines, Red bars: **Sejet** lines **(b)** White clover lines

### Genomic estimations and correlations

#### Faba bean

The narrow-sense heritability at plot level for faba bean grain yield was 0.18, and the broad-sense heritability was 0.23, which indicates that the genetic control of yield is largely due to additive genetic effects and to a smaller extent due to non-additive genetic effects. It is likely, that with larger datasets it would be possible to determine whether such differences are due to non-additive effects or non-captured genomic regions.

On the other hand, the narrow-sense and broad-sense heritabilities for total root length of 0.07 and 0.13, respectively, indicate that additive and non-additive genetic effects were approximately equally important for root length, and that root length could also be strongly affected by other factors, such as environmental effects or GxE interactions.

Based on the bivariate model for grain yield and total root length, the estimated additive genetic correlation between the two traits was 0.83 and the estimated correlation of the line effects was 0.53. So, these two traits seem to be highly genetically correlated, albeit at a quite high level of standard errors in both cases, 0.47 and 0.45 respectively (Table 1, Figure 5). Nevertheless, distribution analyses confirmed the robustness of the estimated genetic correlation between greenhouse total root length (TRL) values and field grain yield: A histogram (Figure 7a) reveals a strongly positive correlation, with the posterior distribution peaking near 0.8 and the majority of the probability mass falling between 0.3 and 1.0. As the 95% Bayesian Credible Interval (CI) does not cover zero, there is strong evidence that the genetic correlation is positive, distinct from zero, and statistically credible, and a trace plot (Figure 7b) confirms that the model has successfully converged with well-mixed and stationary sampling. This suggests that the genetic mechanisms driving early root length are reliably linked to seed yield performance in field conditions, indicating that phenotyping and selection for early root development components, particularly total root length could potentially be useful in breeding programs to increase the genetic gain for grain yield.

**Table 1:** Genomic estimations and heritabilities - Faba bean.

|  | Variance component |  |  |  | Heritability |  |
| --- | --- | --- | --- | --- | --- | --- |
| | Block | $Id_g$ | $Id_l$ | e | $h^2$ | $H^2$ |
| **Grain yield** |  |  |  |  |  |  |
| Estimate (*Standard error*) | 15.2 (1.1) | 8.8 (2.3) | 2.6 (1.4) | 23.1 (0.6) | 0.18 | 0.23 |
| **Protein content** |  |  |  |  |  |  |
| Estimate (*Standard error*) | 0.15 (0.03) | 1.35 (0.26) | 0.16 (0.13) | 1.33 (0.04) | 0.45 | 0.50 |
| **Total root length** |  |  |  |  |  |  |
| Estimate (*Standard error*) | - | 510.1 (473.8) | 405.0 (529.1) | 6356.3 (516.3) | 0.07 | 0.13 |

**Figure 5:**
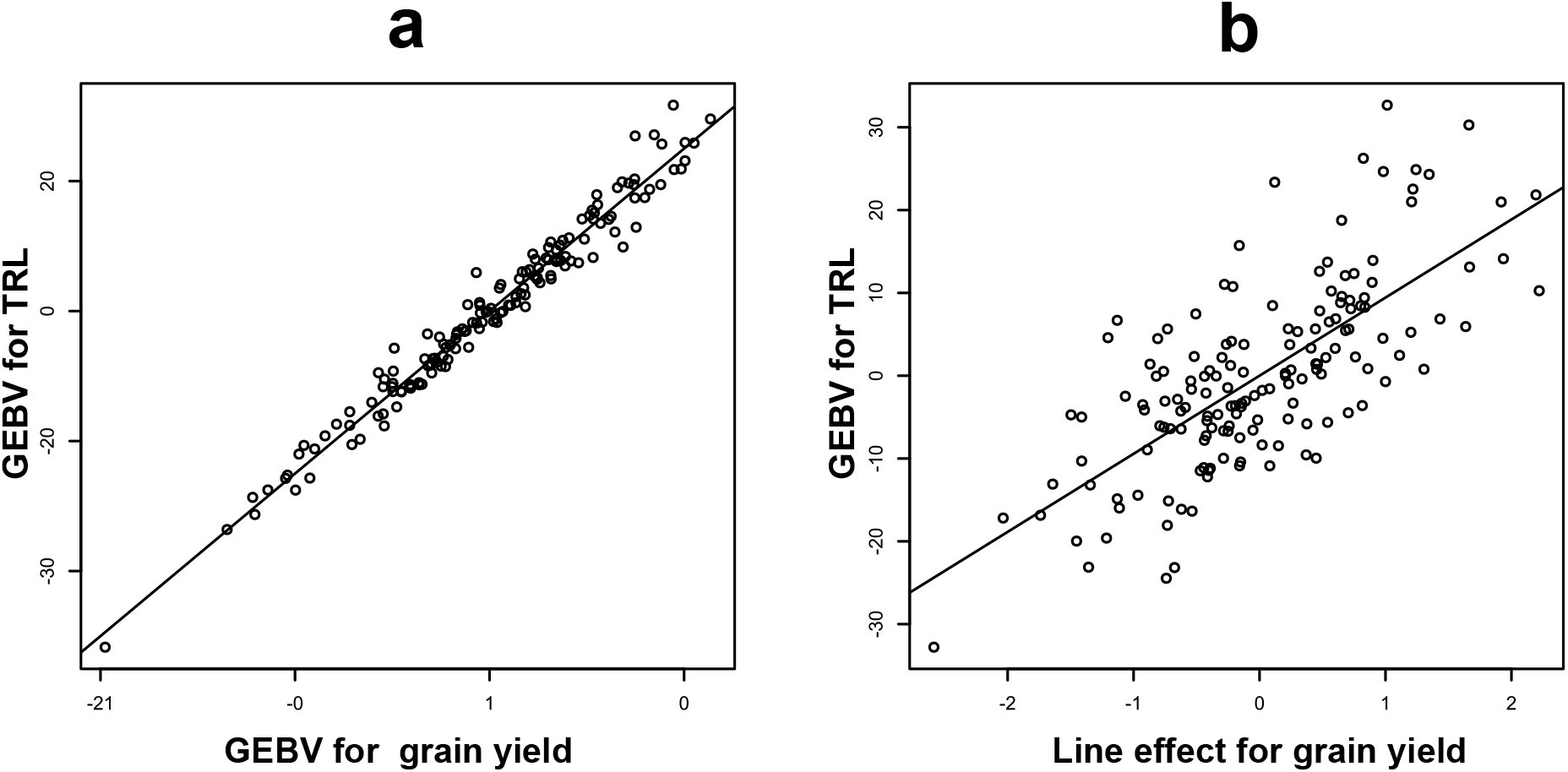
Correlations between Genomic Estimated Breeding Values in faba bean breeding lines. **TRL**: Total root length

#### White clover

We found a positive correlation of 0.17 between GEBVs of total root length (TRL), measured in rhizoboxes and field yield, although the heritability of TRL was very low, which can be explained by the restricted availabilty of field phenotype data, as only a few lines were phenotyped for all traits in two replicates, and therefore the estimates of the variances, breeding values, and correlations could not be very accurate.

On the other hand, phenotyping for leaf parameters could be carried out in the greenhouse for all lines, for which genetic variant data was available. Moderately positive genetic correlations of 0.26 between estimated breeding values of yield from field cuts and of both leaf size and of leaf solidity were identified. For leaf size and leaf solidity, moderate to high narrow-sense heritabilities of 0.46 and 0.63, respectively, were estimated. Both traits had higher narrow-sense heritabilities than 0.22 that was estimated for field yield, and they could therefore be suitable indicator traits in breeding programs for improving yield by indirect selection. The leaf traits can be measured on few, young plants grown in controlled greenhouse conditions, and selection based on these traits might thereby be used at low costs in early stages of breeding programs to increase the genetic gain of yield. Furthermore, phenotyping such traits on selection candidates may improve genomic prediction accuracies for yield in cases where larger training sets are available to use for multivariate models (Table 2, Figure 6).

**Table 2:**
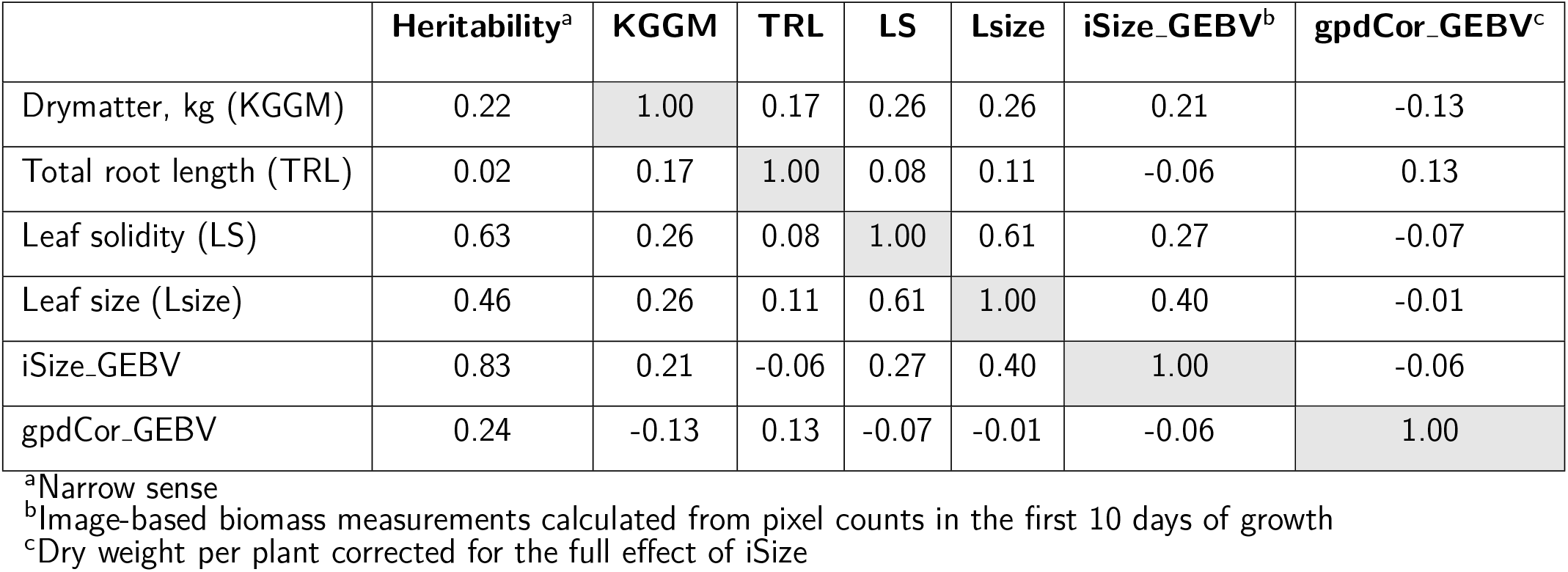
Genomic estimations and heritabilities - White clover.

|  | Heritability<sup>a</sup> | KGGM | TRL | LS | Lsize | iSize_GEBV<sup>b</sup> | gpdCor_GEBV<sup>c</sup> |
| --- | --- | --- | --- | --- | --- | --- | --- |
| Drymatter, kg (KGGM) | 0.22 | 1.00 | 0.17 | 0.26 | 0.26 | 0.21 | -0.13 |
| Total root length (TRL) | 0.02 | 0.17 | 1.00 | 0.08 | 0.11 | -0.06 | 0.13 |
| Leaf solidity (LS) | 0.63 | 0.26 | 0.08 | 1.00 | 0.61 | 0.27 | -0.07 |
| Leaf size (Lsize) | 0.46 | 0.26 | 0.11 | 0.61 | 1.00 | 0.40 | -0.01 |
| iSize_GEBV | 0.83 | 0.21 | -0.06 | 0.27 | 0.40 | 1.00 | -0.06 |
| gpdCor_GEBV | 0.24 | -0.13 | 0.13 | -0.07 | -0.01 | -0.06 | 1.00 |

<sup>a</sup>Narrow sense<sup>b</sup>Image-based biomass measurements calculated from pixel counts in the first 10 days of growth<sup>c</sup>Dry weight per plant corrected for the full effect of iSize

**Figure 6:**
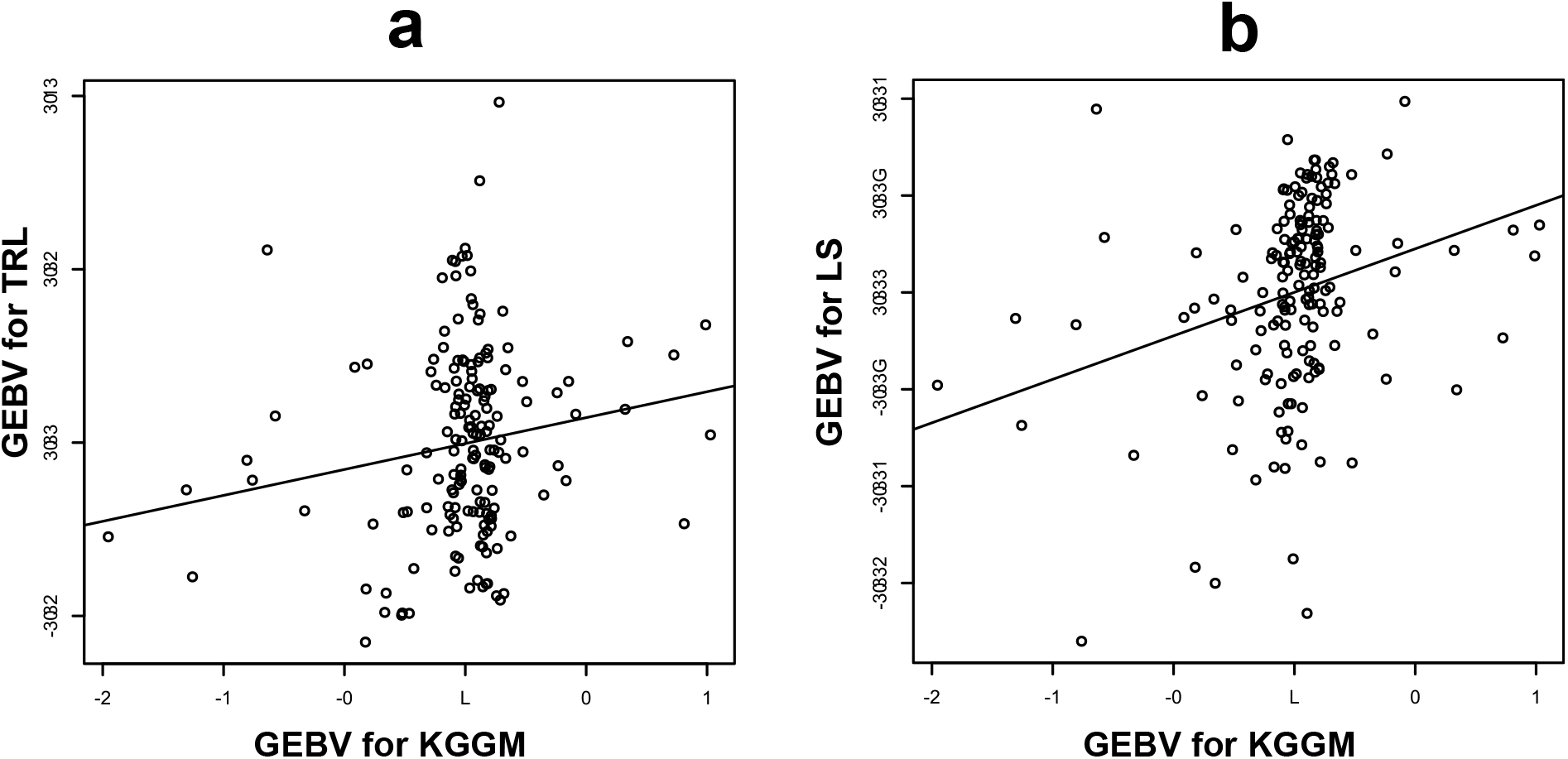
Correlations between Genomic Estimated Breeding Values in white clover breeding lines. **KGGM**: Green matter, kg, **TRL**: Total root length, **LS**: Leaf solidity

**Figure 7:**
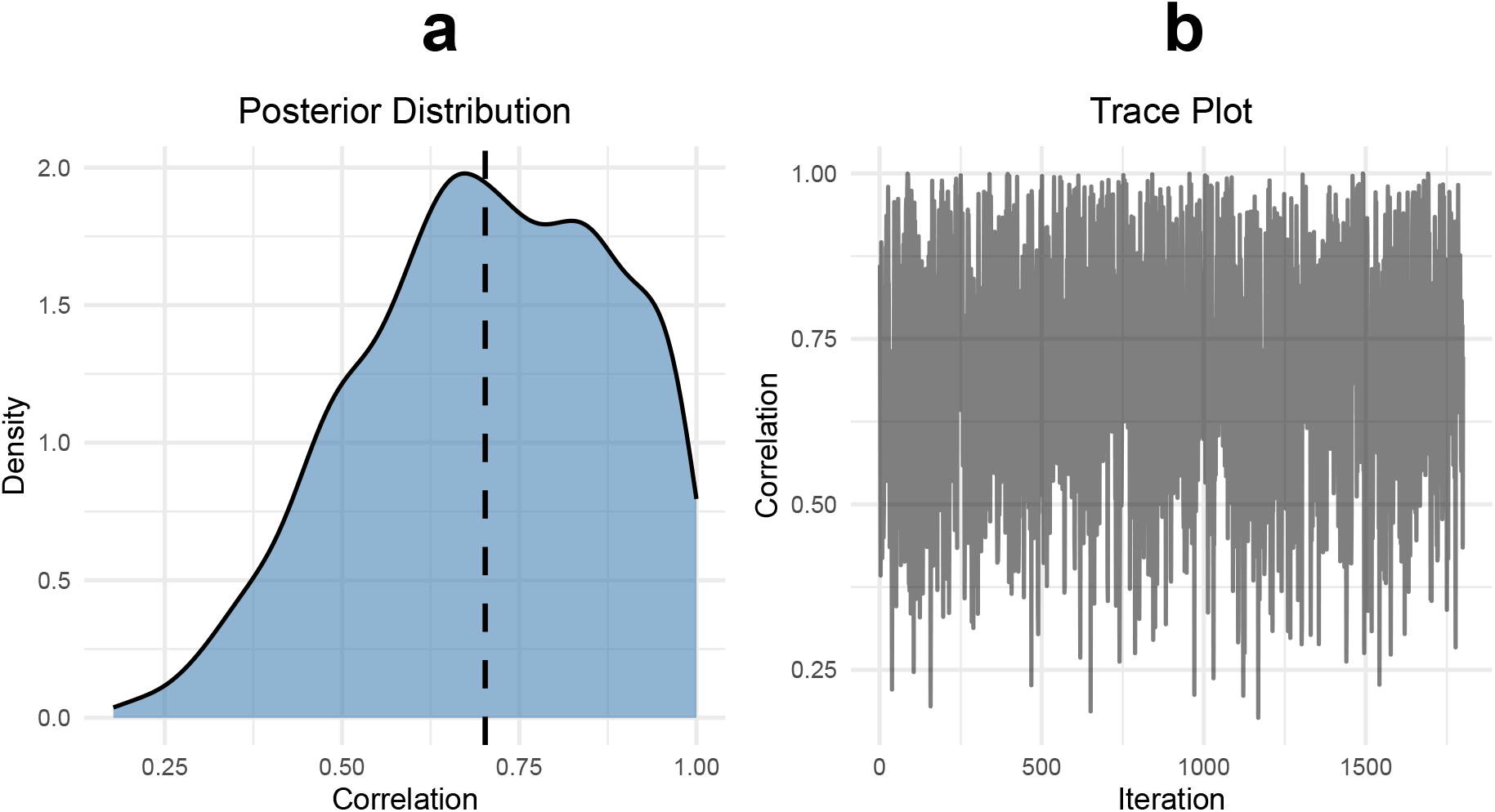
Posterior distribution and trace plot of the genetic correlation between total root length (**TRL**) and grain yield in faba bean. **(a)** Histogram of posterior samples for the correlation parameter between the random intercepts of the two traits **(b)** Trace plot (MCMC chain), indicating the sampling behavior over 3,000 iterations

## Conclusion

Incorporating quantitative phenotype traits that can be assessed under controlled environmental conditions in early developmental phases, while positively correlating with field yield components, can improve the predictability of field performance of crops in different stages of breeding programs.In accordance with earlier findings, our results confirmed that early root development traits can be used as prognostic indicators for field yield across different environments and locations: Both in faba bean as well as in white clover we found significant genetic correlations between Genomic Estimated Breeding Values (GEBVs) of total root length assessed in rhizoboxes and harvested yield in field plot experiments.These findings highlights the value of early root development traits in enhancing genetic gain in breeding programs. Currently, the accessibility to state-of-the-art sequencing technologies and high quality genomic resources effectively support efficient and accurate genomic prediction in all important crop species, but the availability of robust and reliable field phenotype data, well representing the genetic variations targeted by the study, often remains a bottleneck in such projects. Another important aspects of genomic prediction are the availability, development and implementation of appropriate statistical models and methodologies. Our results demonstrated that multi- and bivariate models along with the conceptual pipeline provided in the present case study are well applicable for correlating early root development traits and field yield components. Consequently, these results suggest that early root development traits are of high predictive potential in connection with field yield components, especially if field phenotype data are available to a limited extent.

## Acknowledgements

This work was supported by Promilleafgiftsfonden for Landbrug grant Cultivation of protein crops with a low environmental and climate footprint for the future climate (41647 Klimaprotein). Faba bean genotyping- and field experiment data were generated in the frame of the IMFABA project, supported by The European Union’s Horizon 2020 Programme for Research & Innovation in the ERA-NET Cofund SusCrop project ProFaba (Grant no. 771134). White clover genotying- and greenhouse biomass data were generated in the frame of the NCHAIN project, funded by the Innovation Fund Denmark (Grant no. 4105-00007A).

## Author contributions

Conceptualization: TA, Methodology: IN, MM and PSK, Investigations: IN, LKN, AS and NR, Data analysis: IN, PSK and EB, Writing: IN, PSK and MM, Funding acquisition: TA and SUA

## Supporting Information

### Supplementary material and methods

#### Root image analysis and quantification using RootPainter

##### Image preprocessing

Scanned rhizobox images were trimmed to remove the white frames of the boxes, and images were processed to enhance the contrast between roots and dark background. Then, JPG format files were converted to PNM format applying the following commands of the convert tool of the ImageMagick software:

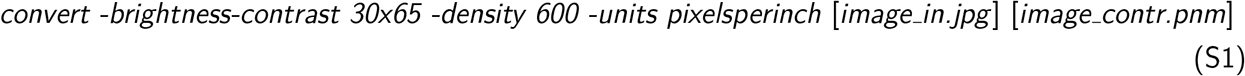

After conversion, a gamma-correction was carried out to restore the original color balance of the images using the pngamma tool of the netpbm software (https://netpbm.sourceforge.net/doc/index.html), and images were converted back to the RootPainter-compatible JPG format:

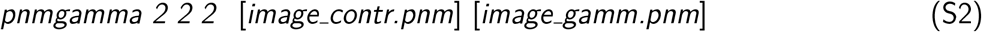

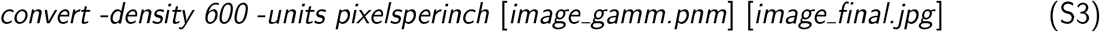

All preprocessing steps were carried out in one go in automatized batch mode using a custom shell script on a Linux workstation.

##### Root segmentation and image-based root length measurement

RootPainter segmentation using convolutional neural network-based deep learning modeling and subsequent analyses were carried out on a local GPU server according to the RootPainter manual (https://github.com/Abe404/root-painter). In brief, after initial manual annotation (marking root-representing clear regions and segments of the dark background by specific colors), a self-learning network training procedure was started on a random set of 30 images with a duration of two to three hours and the resulting intermediate model was saved. After this, an interactive corrective annotation procedure was performed, in which processed images of the selected random set were taken one by one, and annotation errors were manually corrected on them. A gradual annotation accuracy improvement was expected in each subsequent image. If the first training set was exhausted before a desired accuracy level was achieved, the training procedure was repeated using another random set of images. Models resulting from separate training procedures were collected using the “Segment folder” function of the RootPainter software, and they were used individually or in ensembles as input in subsequent training sessions, until a satisfying annotation accuracy level was achieved. The best performing model was used for segmentation of the whole trial image dataset. Segmented images were used to extract Total Root Length (TRL) values for each image, based on the total number of root pixels and expressed as cumulative root length in mm for all plants grown in the same rhizobox. Training and model optimization was carried out separately for the two species (faba bean, white clover) of the study.

**Table S1:** Root features extracted by RhizoVision (Seethepalli et al., 2021)

| Feature extracted | Description |
| --- | --- |
| Surface area ($px^2$, $mm^2$) | *Calculated as the length of the pixel multiplied by the circumference of the cross-section of the root at that pixel* |
| Average diameter ($px$, $mm$) | *The distance transform value at each skeletal pixel is the radius at that pixel and is doubled to give the diameter* |
| Number of branching points ($nr.$) | *Pixels in the identified root topology that have at least three neighboring skeletal root segments* |

##### Image-based analysis using RhizoVision

Segmented root images created by RootPainter were exported as black and white images and further processed by the RhizoVision software. Features as described in Table S1 were extracted using default parameters.

#### Leaf color analysis in white clover

Average pixel RGB values were calculated from the 400px image files by the *convert* tool of the ImageMagick software using a single-line command including the following formula:

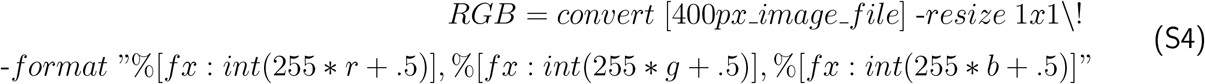

For each line, RGB values were converted to perceived brightness values using the following formula (Luma conversion for perceived brightness, in which the green component has the main effect):

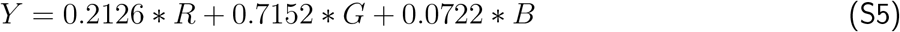

#### Lists of plant genotypes used in the study

**Table S2:** Faba bean lines used in the rhizobox and field experiments.

| Name/ID | Alternative Name | Source of seeds<sup>a</sup> |
| --- | --- | --- |
| **Standard varieties** |  |  |
| Lynx |  | NS |
| Fuego |  | NS |
| Taifun |  | AU |
| Stella |  | NS |
| Bolivia |  | NS |
| Protina |  | AU |
| Capri |  | AU |
| **IMFABA Core lines** |  |  |
| Core25 | Gelber Melaninfleck | NS |
| Core48 | Mythos | NS |
| Core64 | Tiffany | NS |
| Core81 | Stabil | NS |
| Core115 | 11689 | NS |
| Core118 | 13039 | NS |
| Core134 | 124097 | NS |
| Core135 | 124138 | NS |
| Core136 | 124229 | NS |
| Core137 | 124353 | NS |
| Core138 | 130731 | NS |
| Core142 | Aurora | NS |
| Core144 | EH95078-6 | NS |
| Core147 | ILB938/2-2 | NS |
| Core149 | Melodie | NS |
| Core188 | PI377555 | NS |
| Core195 | PI430714 | NS |
| Core239 | PI458603 | NS |
| Core246 | PI469138 | NS |
| Core324 | INK2 | NS |
| Core342 | 687-8 | NS |
| Core401 | NV639-1 | NS |
| Core421 | RIL168L | NS |
| **Breeding lines<sup>b</sup>** |  |  |
| SJ: 75 lines |  | SJ |
| NS: 75 lines |  | NS |

<sup>a</sup>NS: Nordic Seed; AU: Aarhus University; SJ: Sejet
<sup>b</sup>Breeding lines with company-owned codes

**Table S3:** White clover lines used in the rhizobox and field experiments.

| Source variety | Nr. of lines |
| --- | --- |
| AberBoost | 10 |
| AberConcord | 10 |
| AberPearl | 10 |
| Aran | 7 |
| Avalon | 7 |
| Barblanca | 10 |
| Borek | 8 |
| Brianna | 9 |
| Chieftain | 8 |
| Coolfin | 10 |
| Cyma | 10 |
| Iona | 10 |
| Klondike | 6 |
| Liblanc | 7 |
| Liflex | 7 |
| Merida | 10 |
| Rabbani | 9 |
| Riesling | 8 |
| Silvester | 10 |
| Violin | 10 |

## Supplementary figures

**Figure S1:**
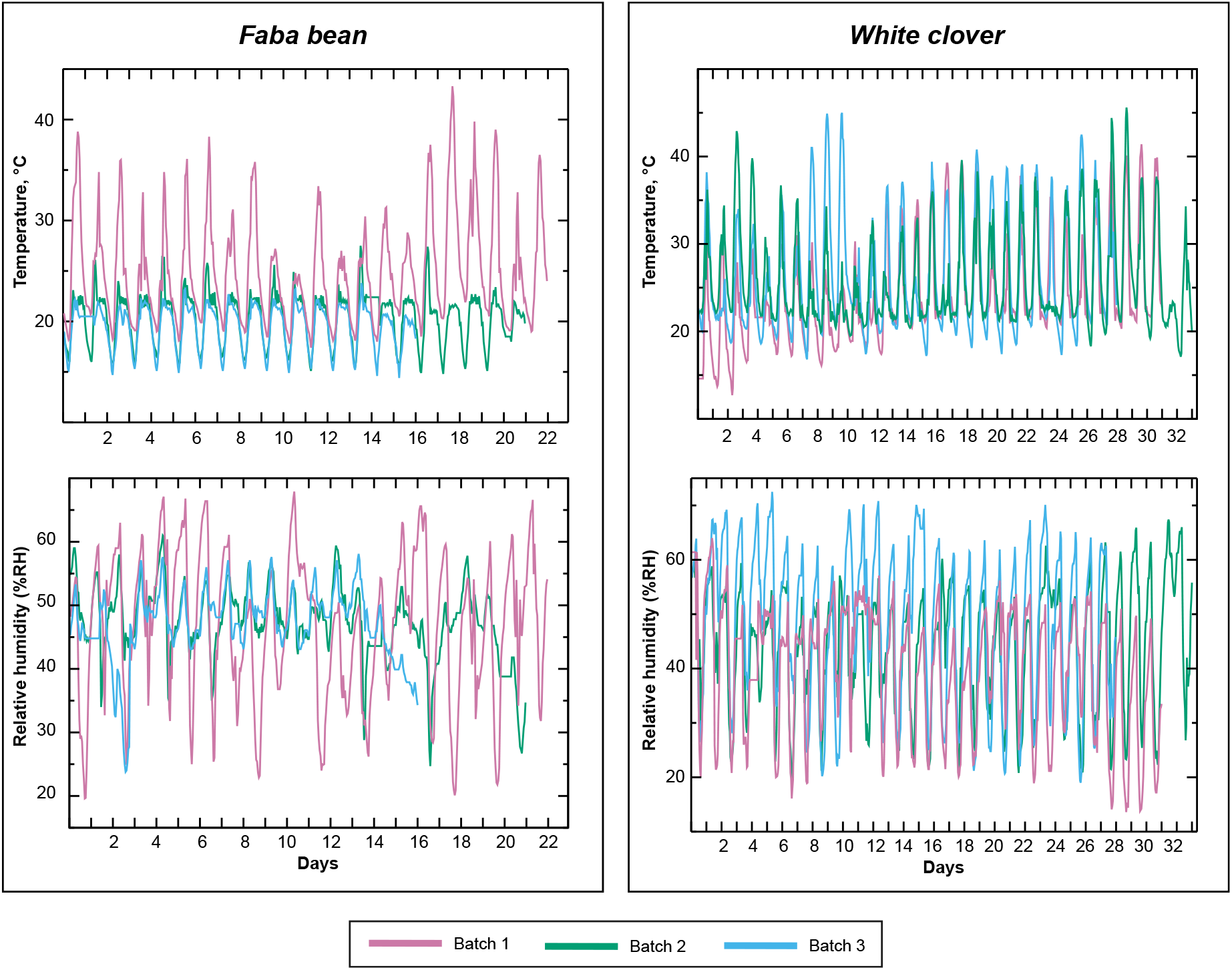
Plots of greenhouse climatic parameters (Temperature, RH) during faba bean and white clover rhizobox experiments

**Figure S2:**
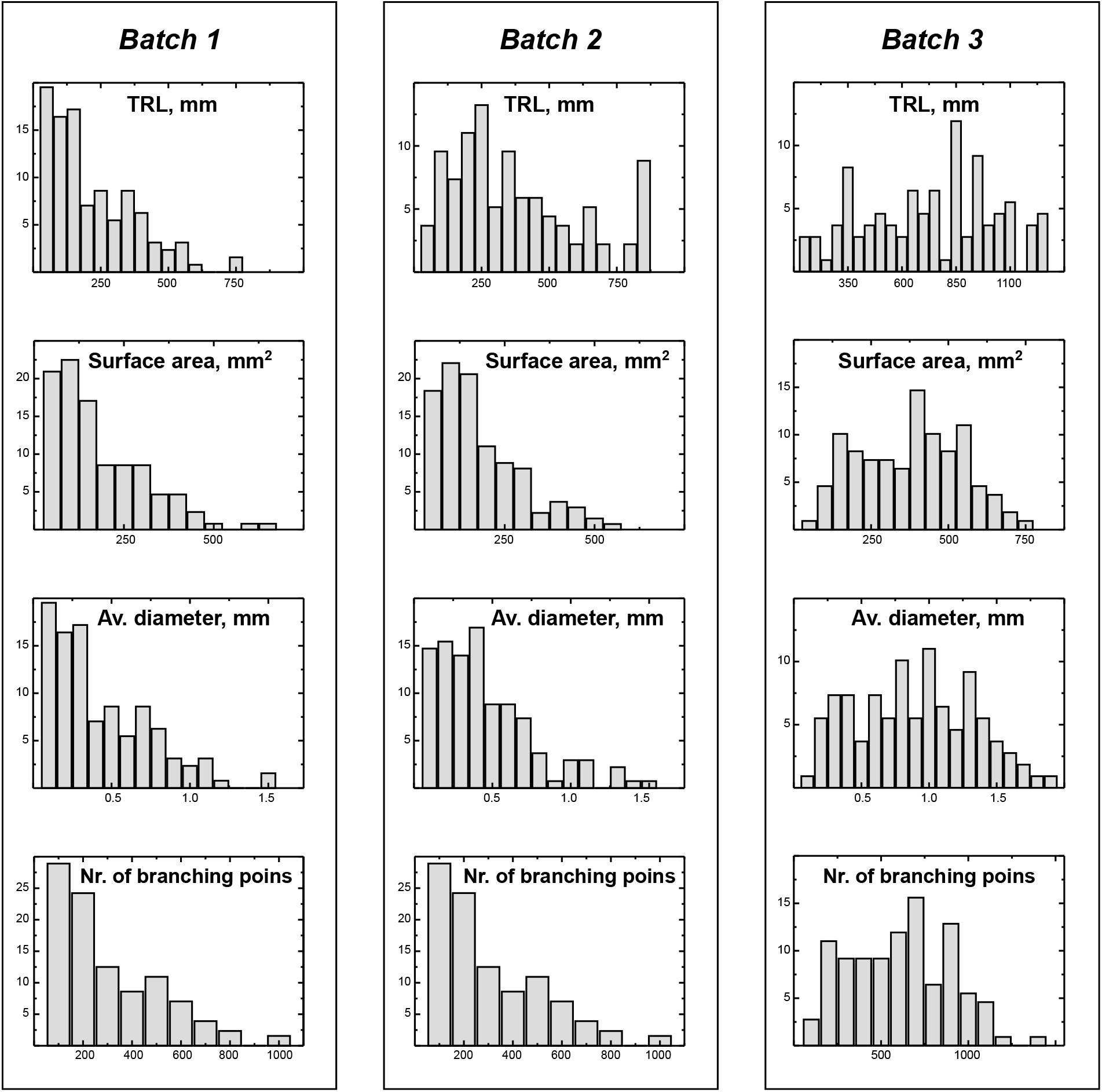
Distribution of quantitative root parameter values in three white clover rhizobox batch experiments

